# Policy precision reveals action-phase impulsivity in women with premenstrual syndrome during risk-taking

**DOI:** 10.64898/2026.03.12.711243

**Authors:** Dayoung Yoon, Bumseok Jeong

**Author notes:** Contributing authors *Email addresses:* (Dayoung Yoon), (Bumseok Jeong).

## Abstract

The Balloon Analogue Risk Task (BART) is widely used to assess risk-taking and impulsivity, yet existing computational models struggle to unify sequential and prior evaluation strategies or fully capture uncertainty-driven information-seeking behavior. To address this, we introduce a novel computational framework grounded in the Active Inference Framework (AIF), which conceptualizes behavior as the minimization of expected free energy. Model comparisons demonstrate that AIF-based models statistically outperform existing benchmarks. Furthermore, we applied this framework to investigate impulsivity in women with Premenstrual Syndrome (PMS). Our model revealed that the PMS group exhibited significantly higher values in inverse precision of policy (*β*_0_) and the phase difference of this parameter was only observed in PMS group. This suggests that high *β*_0_ serves as a robust computational marker, reflecting both the trait impulsivity inherent in PMS and its state-like exacerbation across the menstrual cycle. Lastly, our findings indicate that impulsivity in PMS manifests not as a learning deficit, but as heightened sensitivity to trial-by-trial sequential evaluation at the expense of stable, pre-planned prior policies. This framework provides a neurobiologically plausible and mechanistically granular understanding of risk-taking, offering new avenues for computational psychiatry.

## Introduction

The Balloon Analogue Risk Task (BART) was developed to measure risk-taking behavior and is designed to assess how individuals balance potential rewards against the possibility of loss (Lejuez et al., 2002; van Ravenzwaaij et al., 2011). Behavioral metrics derived from this task have been widely used as indicators of impulsivity. Various computational models have been proposed to explain behavior in BART. For example, (Wallsten et al., 2005) introduced a choice model incorporating two distinct strategies. One is Sequential evaluation and choice, where a participant evaluates the two options—pump or stop—at each pump opportunity. The other is prior evaluation and choice, where a participant decides the optimal number of pumps before starting each trial and acts accordingly. Model comparison showed that the latter strategy yielded better explanatory power on the observed data. However, (Park et al., 2021) introduced the EWMV (Exponential Weighted Mean and Variance) model, which adopts the sequential evaluation approach and demonstrated superior performance. This suggests that even minor modifications in parameterization or utility computation can change which model best fits the data. Thus, the two strategies should not be viewed as mutually exclusive mechanisms; rather, they may reflect different phases of the same underlying decision process. Specifically, when the belief state is highly uncertain—particularly during early trials— participants may rely on the sequential evaluation strategy. As learning progresses and uncertainty diminishes, their behavior may gradually shift toward a prior evaluation mode, such as deciding in advance, “I’ll pump about eight times.” In other words, Sequential evaluation is alternated with prior evaluation as confidence in belief state increases.

In the EWMV model introduced by (Park et al., 2021), the agent performs sequential evaluation by computing the utility of pumping at every pump opportunity. Specifically, the utility of pumping at the h-th pump in trial *t*, 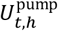, is defined as follows, incorporating not only the mean but also the variance of potential outcomes:

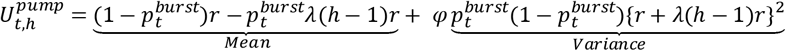

Here, *φ* denotes the risk preference parameter, such that larger values of *φ* increase the weight of the variance term and encourage risk-taking behavior. Hence, risk-seeking individuals are more likely to continue pumping under uncertainty. Since 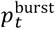 starts from 0.5 in the initial, fully uncertain state and gradually decreases as pump successes are observed more frequently than failures, the variance term decreases as 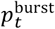 falls below 0.5. In contrast, the mean term increases as 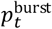 decreases. Because these two terms change in opposite directions depending on uncertainty, the parameter *φ* in principle modulates the balance between them and can therefore explain the transition between sequential evaluation and prior evaluation as confidence in 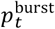 increases.

However, the parameter estimates reported in the original study show that *φ* falls within an extremely narrow range (approximately −0.003 to 0.005). Within such a small interval, the utility monotonically decreases as 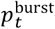 increases, and the contribution of the variance term is effectively negligible. Thus, the model does not meaningfully capture the intended risk sensitivity through the variance component.

Furthermore, increases in loss aversion (*λ*) exert opposing effects on the two components of the utility: greater *λ* decreases the mean utility while simultaneously increasing the variance term. While it is intuitively consistent that stronger loss aversion should reduce the tendency to pump, increasing the variance—and thereby increasing pump tendency—produces a counterintuitive effect within the model. If *φ* were negative, the increased variance would reduce the utility of pumping, but in that case the model would no longer be able to explain exploratory pumping behavior driven by greater uncertainty (i.e., increased variance when 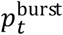 is uncertain).

These findings highlight two conceptual limitations of existing models: first, the need for a unified approach that integrates rather than separates the two choice strategies, and second, the need for a framework that explicitly incorporates the influence of belief uncertainty on decision-making. To address both issues simultaneously, we propose a new model based on the Active inference framework (AIF).

In AIF, agents make decisions by minimizing the expected free energy (EFE), which naturally combines both pragmatic and epistemic components. The pragmatic value corresponds to the expected utility term in the EWMV model, whereas the epistemic value represents the information gain associated with uncertainty reduction. When *p*^*burst*^ is uncertain, an increased tendency to pump can therefore be interpreted as an information-seeking behavior driven by the epistemic component. Unlike traditional models that linearly combine mean and variance with separate weighting parameters, AIF computes both components directly from the underlying belief distributions. Specifically, if *q*(*θ*) denotes the belief distribution over states and *p(o)* the preference distribution, the EFE (*G*) can be expressed as

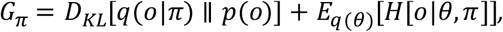

where the first term represents pragmatic value and the second corresponds to epistemic value. Minimizing EFE thus balances the pursuit of rewards with the reduction of uncertainty in a single computational principle.

Before each trial, the agent evaluates multiple possible policies—for example, pumping once, twice, or up to k times—and computes the EFE for each policy. The policy that minimizes EFE is selected according to

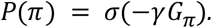

As cumulative rewards increase within a single trial, the potential loss associated with additional pump grows accordingly, thereby shifting the posterior belief distribution in real-time. This dynamic shift often leads to deviations from the initially planned number of pumps. For instance, a participant who originally intended to pump eight times may prematurely stop at the sixth pump to secure accumulated rewards (loss aversion) or, conversely, may opt for additional pumps if their updated confidence suggests a higher threshold for bursting.

Over repeated trials, as the estimation of *q*(*θ*) becomes more accurate, the discrepancy between the initially planned number of pumps and the actual performance in a single trial should systematically decrease. We validated this foundational premise of our model by demonstrating that the difference between the EFE-minimizing prior policy and actual choice significantly diminishes from early to late task stages (Figure 1A). Furthermore, the simulation results demonstrate that as the estimation of the bursting point becomes more precise over repeated trials, the standard deviation in the number of pumps decreases, leading to a more stabilized pumping strategy (Figure 1B, C). The existence of such behavioral deviations and their subsequent stabilization suggest a dual-process mechanism in decision-making: an initial pre-planned strategy (prior evaluation) is continuously modulated by a pump-by-pump re-evaluation (sequential evaluation).

**Figure 1.**
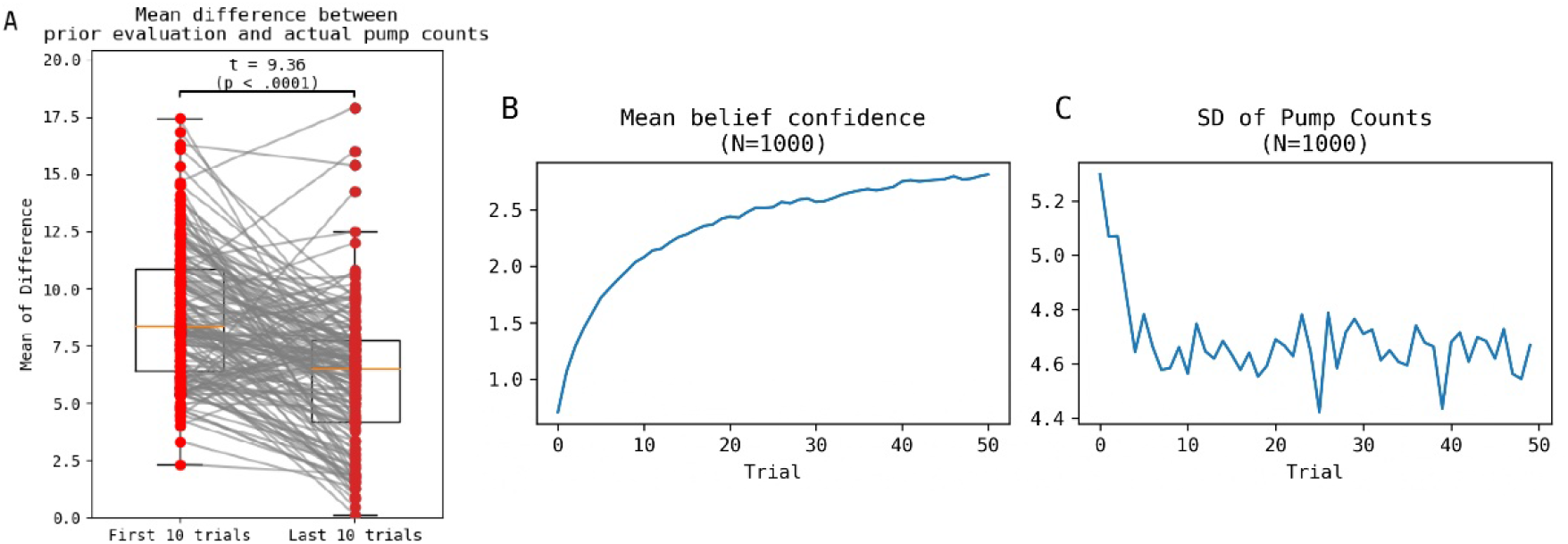
(A) Boxplots illustrate the average difference between the pre-planned pump counts (minimizing expected free energy before each trial) and the actual pumps performed by participants (N=159). Individual data points are connected by lines to show the longitudinal shift. The significant reduction in this discrepancy from the initial 10 trials to the final 10 trials demonstrates that subjects’ sequential actions increasingly align with their prior evaluative strategies as they gain information. (B-C) Results from 1000 simulated Active inference agents show (B) the trajectory of belief confidence regarding the balloon’s bursting point and (C) the standard deviation (SD) of pump counts across simulated agents at each trial. As the estimation of the bursting distribution becomes more precise over repeated trials, belief confidence rises, leading to a marked stabilization of the pumping strategy (decreased SD).

### Learning model

Traditionally, learning models in the BART have estimated the bursting probability through a continuous ratio, such as the number of pops divided by the total number of pumps (Eq. (8)). However, we posit that human agents do not meticulously count every cumulative pump to update their beliefs in real-time. It is more psychologically plausible that agents learn through point-specific heuristics—such as “I have succeeded at the 7^th^ pump most of the time” or “After the 12^th^ pump, the success rate drops to half”—rather than maintaining a meticulous tally of the total cumulative pumps. This suggests that agents estimate the probability of bursting at each pump step individually (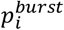, *i*=1, …mp). Then we can calculate the probability of survival at each pump step *i* at the point of the *h*-th pump as follows:

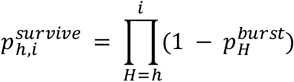

While the probability of survival can be mathematically derived as 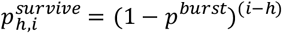, such indirect continuous models fail to capture the discrete, step-like nature of the belief states required for specific task environment, such as those with a fixed bursting point. For instance, if a balloon is programmed to invariably burst at the 10^th^ pump, a human agent, after several trials of observation, would eventually infer this deterministic rule and consistently stop at almost exactly the 9^th^ pump to maximize accumulated rewards and avoid pop. Our simulations revealed that learning models based on the conventional belief state of *p*^*burst*^ were unable to replicate this behavioral pattern (Supplementary Figure 1). By modeling the learning process at each discrete pump opportunity 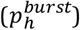, we can more accurately represent the actual cognitive trajectory of human decision-makers in the BART task.

Each pump opportunity corresponds to the Bernoulli trial with outcomes of success or failure of pump, and state value *q*(*θ*_h_) can be represented by a beta distribution, where *θ*_h_ is the probability of the success 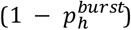.

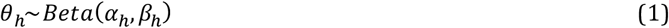

Where *α* and *β* are equal to one plus the number of success or failure outcomes at *h*-th pump, respectively. In this case, as the task progresses, the learning process can be represented simply by adding one according to the outcome of each trial (*o*_*t,h*_), as the likelihood of outcome at each trial is a binomial distribution, which is conjugated to the beta distribution.

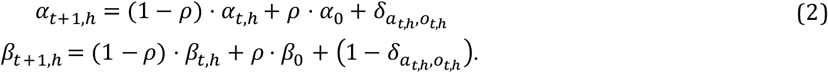

*δ* indicates the Kronecker delta and takes 1 if both variables *a*_*t,h*_ and *o*_*t, h*_ are equal (when a pump action results in success), and 0 otherwise. *ρ* is the decay parameter (0 < *ρ* < 1). Larger value leads the concentration parameters to converge toward their initial values, effectively corresponding to a forgetting process in which learning from observations fades over time. This parameter can be dynamically updated across trials, quantified as predictive surprise which is the gap between prior beliefs and observed outcomes (Liakoni et al., 2021) as follows:

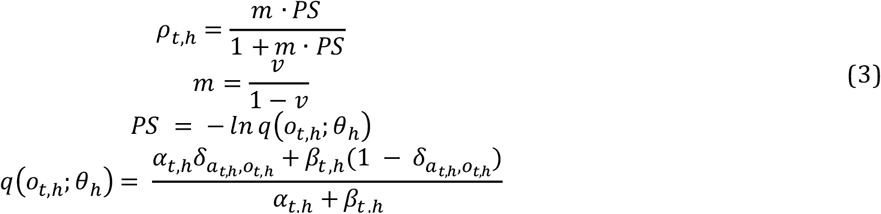

### Action model

Formally, we express the EFE of a policy *π* on pump opportunity *h* on trial *t* as:

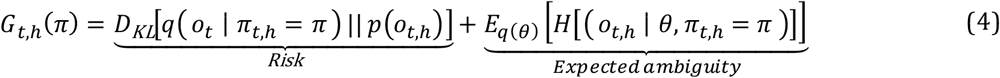

At the beginning of each trial (*h* = 0), the agent considers a set of possible policies corresponding to the maximum number of pumps (*mp*) plus one. These policies represent all possible stopping points: stopping after 0 pumps, after 1 pump, and so on, up to stopping after *mp* pumps. At each pump opportunity, the number of remaining possible policies decreases by one, as one additional pump action has been executed.

Similarly, the sample space of possible outcomes consists of twice the number of these policies, i.e., 2 × (*mp* + 1) elements. Each outcome represents either bursting or stopping after a given number of pumps: bursting after 0 pumps (theoretically included in the space but with zero probability), stopping after 0 pumps, bursting after 1 pump, stopping after 1 pump, and so forth, up to bursting after *mp* pumps and stopping after *mp* pumps. At each pump opportunity, two outcomes are removed from this space, corresponding to the updated possible results of bursting or stopping at the current step.

The preference distribution of outcomes, *p*(*o*_*t*,h_), is encoded using *φ* parameters for each participant. The loss associated with a bursting outcome depends only on the accumulated reward up to that point and therefore remains constant across all pump levels, whereas the potential gain increases linearly with the number of pumps. The loss or gain of the possible outcomes at the point of the *h*-th pump in trial *t* can be expressed as follows and subsequently converted into a probability distribution using the softmax rule:

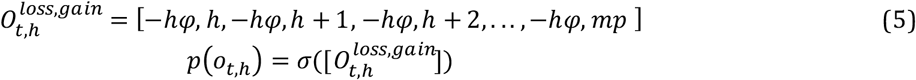

According to the prospect theory (Kahneman & Tversky, 2018), the subjective utility of losses is greater in magnitude than that of equivalent gains. Therefore, a risk aversiveness parameter is multiplied to the loss outcomes to account for this asymmetry (*φ* > 1).

The likelihood of outcomes given a particular policy *k* (stopping after *k* pumps) and state, i.e., under the belief state (*s*), is therefore defined as follows:

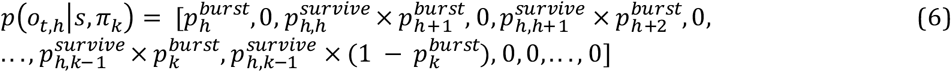

This ensures that the predicted outcome at a future step *k* accounts for the prerequisite that the balloon must not have burst in preceding steps.

The second term in Eq. (4), expected ambiguity, decreases as the policy involves a greater number of pumps. Therefore, at each pump opportunity, the stop policy does not reduce ambiguity any further, resulting in this term being zero (Figure 2C and red line in 2F). This reduction is larger when the variance of ***q***(***θ***) is high, indicating greater uncertainty in the belief about the bursting probability. However, policies with a higher number of pumps also carry an increased probability of failure (Figure 2B), as described in Eq. (6), thereby increasing the overall EFE (Figure 2A). Consequently, the balance between these two opposing terms leads the agent to select an optimal policy.

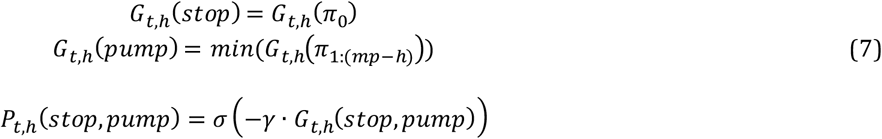

**Figure 2.**
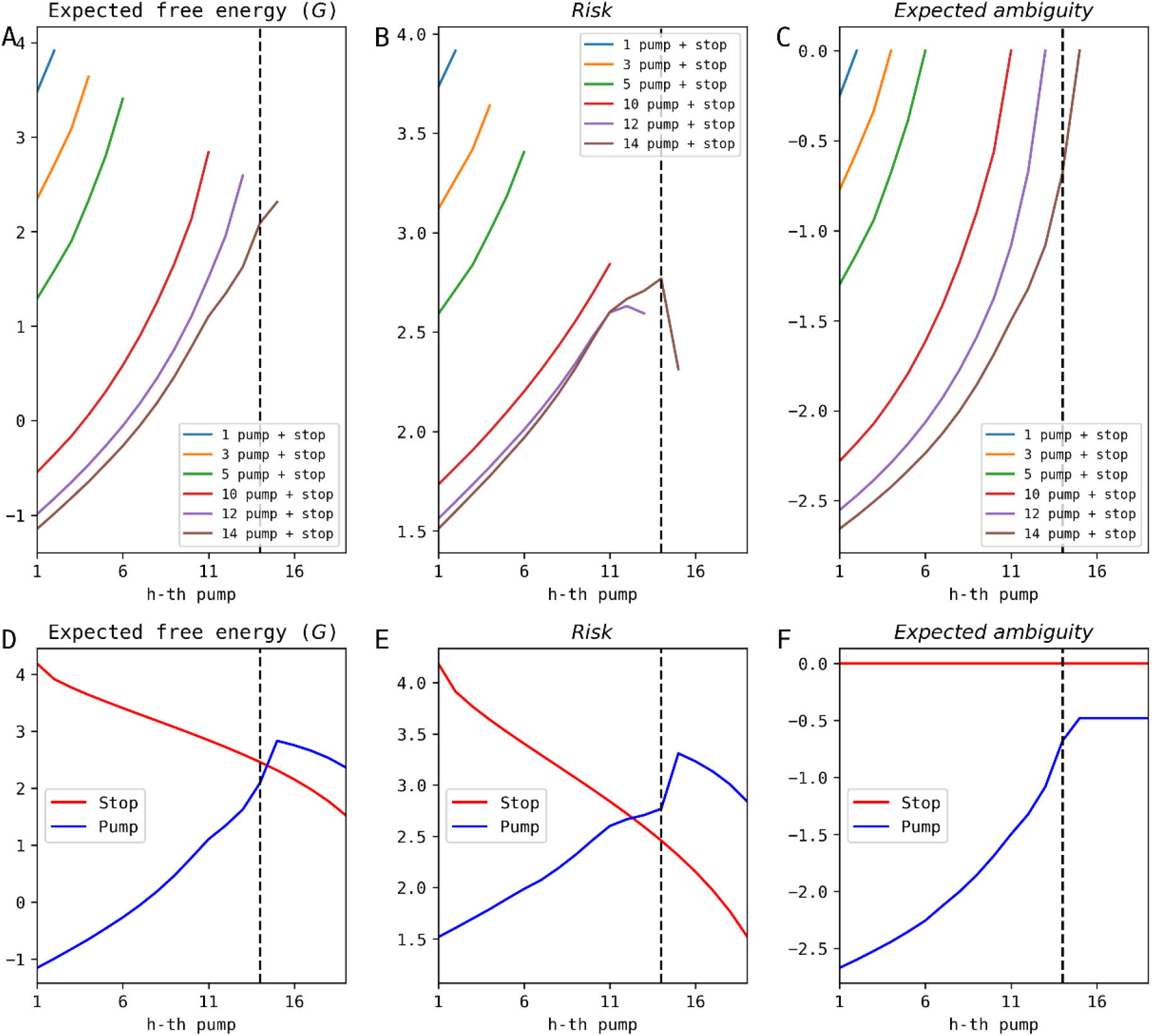
Example plot of expected free energy (EFE) for different policy options at each pump opportunity. Panels show the overall Expected free energy (*G*) (A, D), risk (B, E), and expected ambiguity (C, F). The blue lines in (D-F) show the pump action selected as the minimum *G* at each pump opportunity. The dotted lines mark the point where the *G* of two actions intersect.

The policy precision parameter, ***γ***, can be updated, and it provided a better explanation of behavioral data in the 2-armed bandit task compared to a baseline model with fixed policy precision in our previous study (Yoon et al., 2025). In that study, increases or decreases in ***γ*** modulated the balance between model-based decision-making and influence of previous outcomes, thereby capturing impulsive responses driven by recent feedback. In the current BART task, changes in ***γ*** account for the phenomenon where individuals exhibit heightened engagement under uncertainty, which in turn promotes impulsive behavior such as gambling-like or risk-seeking choices (Fiorillo et al., 2003; Clark et al., 2009; Linnet et al., 2011).

### Impulsivity in Premenstrual syndrome

Women with premenstrual syndrome (PMS) have been shown to report significantly higher levels of behavioral impulsivity and greater difficulties in emotion regulation (Dawson et al., 2018; Ducasse et al., 2016; Petersen et al., 2016; Ko et al., 2014). However, most of these findings rely primarily on self-report questionnaires, and relatively few studies have examined objective behavioral indicators of impulsivity in this population. A small number of studies have investigated cognitive deficits in PMS using laboratory-based cognitive tasks (Bannbers et al., 2012; Yen et al., 2012, 2023). For instance, (Yen et al., 2023) reported that women with PMDD showed significantly poorer response inhibition than controls across both the early and late luteal phases. In contrast, (Bannbers et al., 2012) found no significant difference between PMDD participants and healthy controls in the number of correct or incorrect responses, nor in reaction times, during either the follicular or luteal phases. This substantial inconsistency in overt behavioral outcomes suggests that Go/No-Go performance may lack the sensitivity and reliability required to consistently capture impulsivity-related deficits in PMS. The mixed findings highlight the need for more sensitive, process-based behavioral measures that can capture the underlying cognitive components of impulsivity beyond what is observable from simple accuracy or reaction-time metrics.

A substantial body of prior research has investigated impulsivity using the BART (Acheson et al., 2007; Hunt et al., 2005; Lauriola et al., 2014; Rieser et al., 2019), and (Meda et al., 2009) conceptualized impulsivity as a multidimensional construct and examined five underlying components, reporting that the risk-taking tendency captured by BART represents an independent dimension that is distinguishable from the other impulsivity factors. In the present study, we sought to compare several widely used behavioral indices across groups and task phases, including the adjusted mean pumps (AMP)—the most commonly employed BART measure (Lejuez et al., 2002)—as well as the adjusted variance of pumps (AVP) and post-failure mean pumps (PFMP), which have also been used as behavioral markers of impulsivity (Ashenhurst et al., 2011; Jentsch et al., 2010).

AMP and AVP correspond to the mean and variance of pump counts on balloons that did not explode. (Ashenhurst et al., 2011) proposed quantifying pump variability by dividing the variance by AMP, since variance naturally scales with the average (AVPAMP). PFMP represents the mean number of pumps on trials immediately following a trial failure (i.e., balloon explosion). Previous studies have treated these three measures as behavioral markers reflecting a general risk-taking component of impulsivity. By examining their associations with the computational model parameters, we aim to clarify which specific cognitive processes or latent traits each behavioral index may represent.

## Materials and Methods

### Participants

A total of 159 female participants were recruited (mean age = 26.2 ± 6.6 years). Based on each participant’s reported date of the last menstrual period and average cycle length, participants were categorized as being in either the follicular or luteal phase. Four participants were excluded from the phase-related analyses due to inaccurate reporting of their menstrual information, which made phase classification impossible.

The Premenstrual Symptoms Screening Tool (PSST; Steiner et al., 2003) was used to classify participants into the PMS and control groups. Sixty-six participants were identified as having PMS, and among them, fourteen met criteria for PMDD. In the PMS group, 36 participants were in the luteal phase, while 46 participants in the control group were in the luteal phase. All participants completed the BART online.

The study design was approved by the Institutional Review Board of KAIST (IRB No KH2022-043). All participants provided written informed consent before participating in this study in accordance with the Declaration of Helsinki.

### Behavioral model

We fitted four behavioral model to the data and compared. First model is the model identified as the best-performing model in (Wallsten et al., 2005). In this model, the burst probability *p*^*burst*^ is updated according to

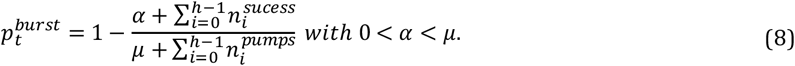

Here, the term 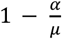 reflects the agent’s prior belief about *p*^*burst*^. This expression corresponds to the posterior expectation of a Beta–Bernoulli Bayesian update based on a prior distribution *p*^*burst*^∼*Beta*(*α, μ* − *α*). Because *p*^*burst*^ is defined as 1 − *E*[*p*^*success*^], the update rule is mathematically equivalent to a standard Beta-distribution update. In effect, this procedure updates the parameters of the Beta distribution in Eq. (2) as:

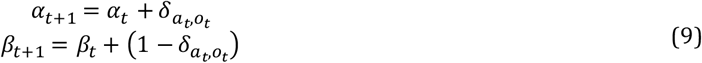

The utility of possible outcomes at the h-th pump on trial *t* is computed, and the pump count that maximizes this utility defines the pump opportunity at which the probability of pumping is 0.5 (Eq. (10)).

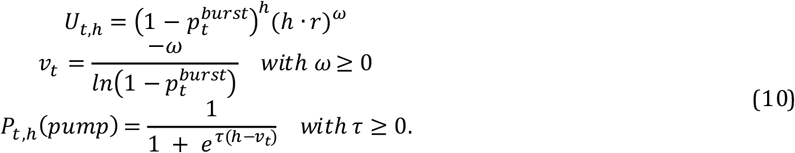

The value *v*_*t*_ is the optimal number of pumps that maximizes *U*_*t*,h_, corresponding to the value of h at which the first derivative of the utility function becomes zero. Based on *v*_*t*_, probability of pump is then computed via a logistic function. Here, *ω* is risk-taking propensity: larger *ω* yields smaller *v*_*t*_, thereby increasing the probability of pumping. The parameter *τ* is the decision temperature, with larger values producing more deterministic choices. Second one is EWMV model.

The other two models are based on the AIF that utilizes the EFE from Eq. (4)-(7). Additionally, the precision of beliefs about actions *γ* in Eq. (7) can be inferred instead of a fixed value (Friston et al., 2014). The initial value of *γ* is calculated from the fixed prior value of *γ*_0_. Thus, the probability that the agent will choose each policy *π*_0_ will be

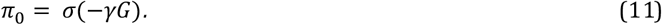

The probability of selecting each policy is determined by its EFE, weighted by precision. Precision is dynamically adjusted based on the alignment between the EFE and the variational free energy (VFE). The VFE (*F*) corresponds to prediction error which is computed as the discrepancy between an agent’s internal model and the observation. Specifically, *F* is computed as the divergence between the prior predictive probability of the balloon reaching a certain size at the *j*-th pump opportunity under policy *π*_*k*_ (i.e., the probability of avoiding an explozion up to that state) and the joint probability after observing the outcome of each pump. Here, *j* indexes the time points (*τ*) in (Smith et al., 2022) about which the agent holds beliefs. The index *j* extends up to (*mp* + 1), representing the maximum time step within the agent’s policy horizon. Specifically, (*j* = *mp* + 1) marks the completion point for the policy of pumping *mp* times and then stopping (*π*_*mp*_). The evaluation is aligned with the time point of the pump policy that minimizes the *G* in Eq. (7). This alignment is necessary because the temporal horizon of the pump policy consistently exceeds that of the stop policy.

Typically, the posterior belief is derived by updating the prior belief to minimize this divergence. However, to ensure computational tractability and to leverage the properties of the beta distribution used to represent the agent’s beliefs, we employ a standard Bayesian update. We assume that the resulting posterior belief represents the optimal distribution that minimizes *F*.

Following the outcome of the (h − 1)-th pump, the agent updates its belief regarding the burst probability distribution 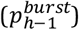. The VFE calculated during this update process serves to modulate the *γ* for the subsequent decision at time step h. We define the minimized *F*_h_ as the divergence between the updated posterior and the prior predictive distribution formed at step h − 1 regarding the balloon’s state at the final time point *j* = *mp* + 1). Under the pump policy (*π*_*k*_) selected in Eq. (**7**)—which minimizes the *G* by planning to stop after *k* pumps—the probability distribution of the balloon’s state remains constant for all *j* > *k*. Specifically, the survival probability is fixed at 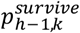, resulting in a state distribution at *j* = *mp* + 1 of:

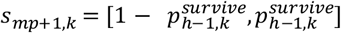

Conversely, the stop policy at step h corresponds to the decision to cease pumping exactly at the h-th opportunity. Under this policy, the state distribution at *j* = *mp* + 1 is represented as:

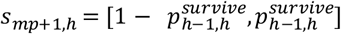

Letting ŝ denote the prior belief, the VFE governing the transition can be formulated as follows:

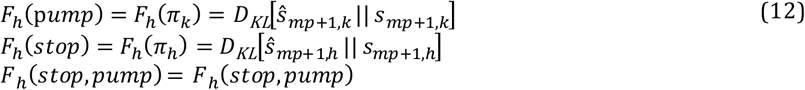

Subsequently, the probability of choosing each policy after a subsequent observation is:

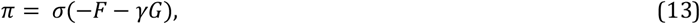

where *F* is calculated relative to the stop policy and the pump policy adopted in Eq. (7). and the error of the agent’s EFE (*G*_*error*_) is given as:

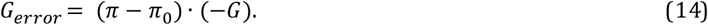

The above equation reflects the level of (dis)agreement between the EFE (G and *π*_0_) and the VFE of the observation (F and *π*) (Smith et al., 2022). Thus, the *γ* is updated based on *G*_*error*_.

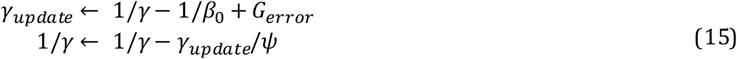

Therefore, policy precision reflects the reliability of EFE in light of newly acquired observations. When the outcome of a selected action violates prior expectations, policy precision decreases, reducing the extent to which action selection is driven by EFE. In the BART, as long as pumping is successful, the VFE of the pump remains lower than that of the stop (Figure 3C).

**Figure 3.**
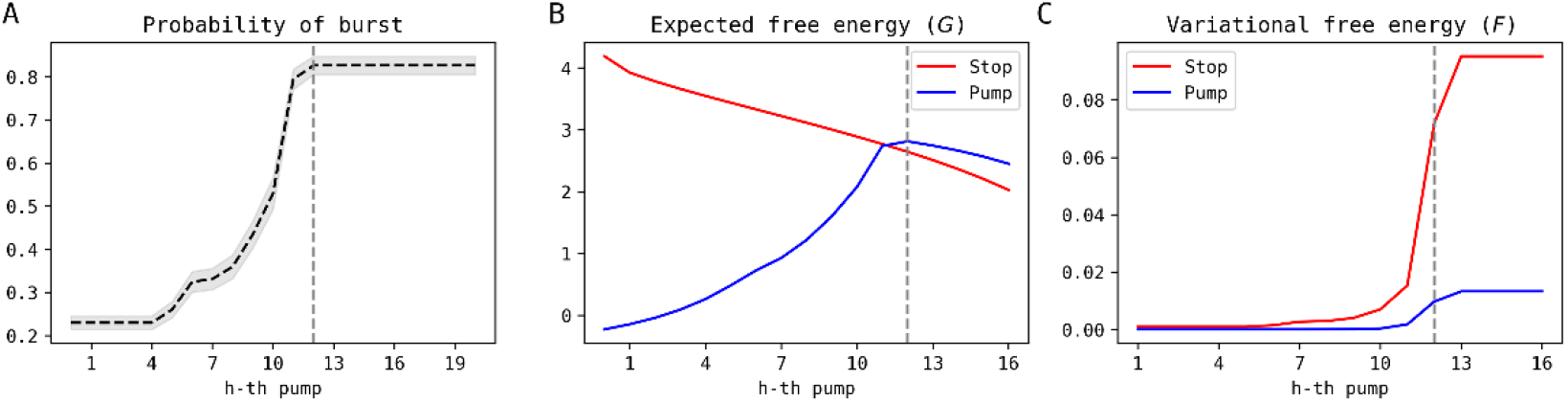
Example plot of probability of bursting at each pump opportunity (A) in the middle of a trial. Based on this belief, a simulation was performed under the manipulation that no explosion occurs until the agent chooses to stop. (B) shows the expected free energy (EFE) for the stop and pump policies at each pump opportunity, and (C) shows the variational free energy (VFE). The VFE of the pump policy remains lower than that of the stop policy as long as pumping is successful. The black dotted lines indicate the point at which the EFE values of the two actions intersect (12 in this example). This point corresponds to the stage where the belief that the balloon will be bursting increases sharply in (A). At this point, the VFE of the stop policy increases steeply, resulting in a larger difference between the two actions.

Consequently, before the EFE of the stop and pump policies intersect, the direction of VFE is aligned with that of EFE, such that increasing policy precision amplifies the influence of EFE on policy selection. In contrast, after the point at which the EFE curves intersect and the EFE of the pump policy becomes higher than that of the stop policy (After the dashed line in Figure 3), the directions of EFE and VFE diverge. Under this condition, a decrease in policy precision counteracts the influence of EFE, thereby suppressing the tendency to stop. As a result of these opposing effects on the pump and stop policies, a sequence of successful pumps acts as a driving force that promotes continued pumping behavior. The influence of *F* increases as the value of *β*_0_ in Eq. (15) becomes larger.

We refer to this model as the *efegc* model (where “*gc*” denotes gamma change) and the model without this step as the *efeX* model as in our previous work (Yoon et al., 2025). We compared four models.

The parameter compositions and recovery results of each model are described in Supplementary Figure 3. Parameter estimation was accomplished using the variational Bayes method (Friston et al., 2007), and the approximations of log model evidence were subjected to random-effect Bayesian model selection (Rigoux et al., 2014).

### Statistical Analysis

To examine the effects of *phase* (pre vs. post) and *group* on each BART-related metric (AMP, VARAMP, PFMP), we conducted a two-way analysis of variance (two-way ANOVA) with interaction terms. For each dependent variable, the model was specified as:

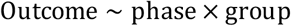

When the ANOVA indicated significant main effects or interactions, post hoc pairwise comparisons were performed using Tukey’s Honestly Significant Difference (HSD) test with familywise error rate controlled at α = .05. For each comparison, mean differences and 95% confidence intervals were estimated. All statistical analyses were conducted in Python using the statsmodels package (ols, anova_lm, pairwise_tukeyhsd).

## Results

### Model Comparison and Comparison of model parameters and BART metrics across group and menstrual phase

The *efegc* model outperformed in four models (Figure 4).

**Figure 4.**
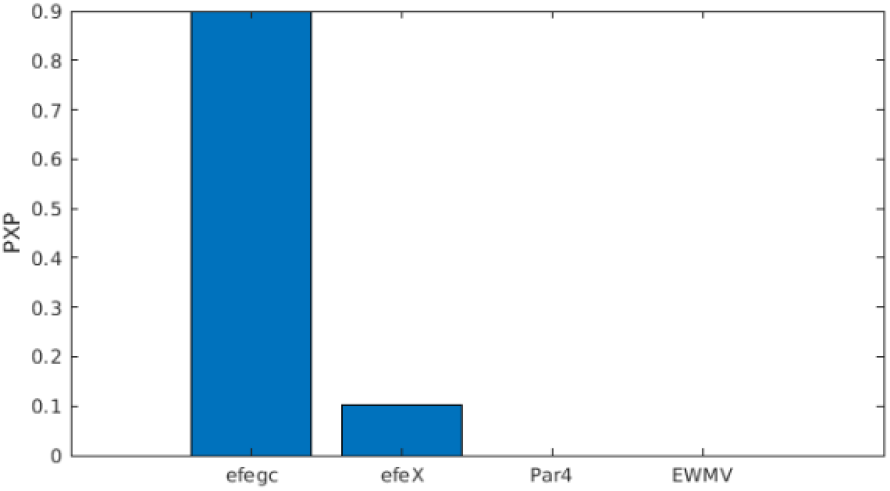
Bayesian model comparison results. (A) The bar plot shows that the proposed active inference model with dynamic policy precision (*efegc*) substantially outperforms the other models in explaining the behavioral data. A higher protected exceedance probability (pxp) indicates stronger evidence in favor of a model.

For AMP and AVPAMP, there were no significant main effects of phase or group, and the phase × group interaction was also not significant (Figure 5A). For PFMP, there was a significant main effect of phase (*F*(1,151)=4.76, *p*=.031), indicating slightly higher values in the follicular phase, as well as a significant interaction (*F*(1,151)=5.08, *p*=.026). To further examine this interaction, we conducted separate comparisons within each group. The results showed no significant phase difference in the PMS group (*t* = 0.20, *p* = .768), whereas a significant phase difference was observed in the control group (*t* = −3.25, *p* = .002).

**Figure 5.**
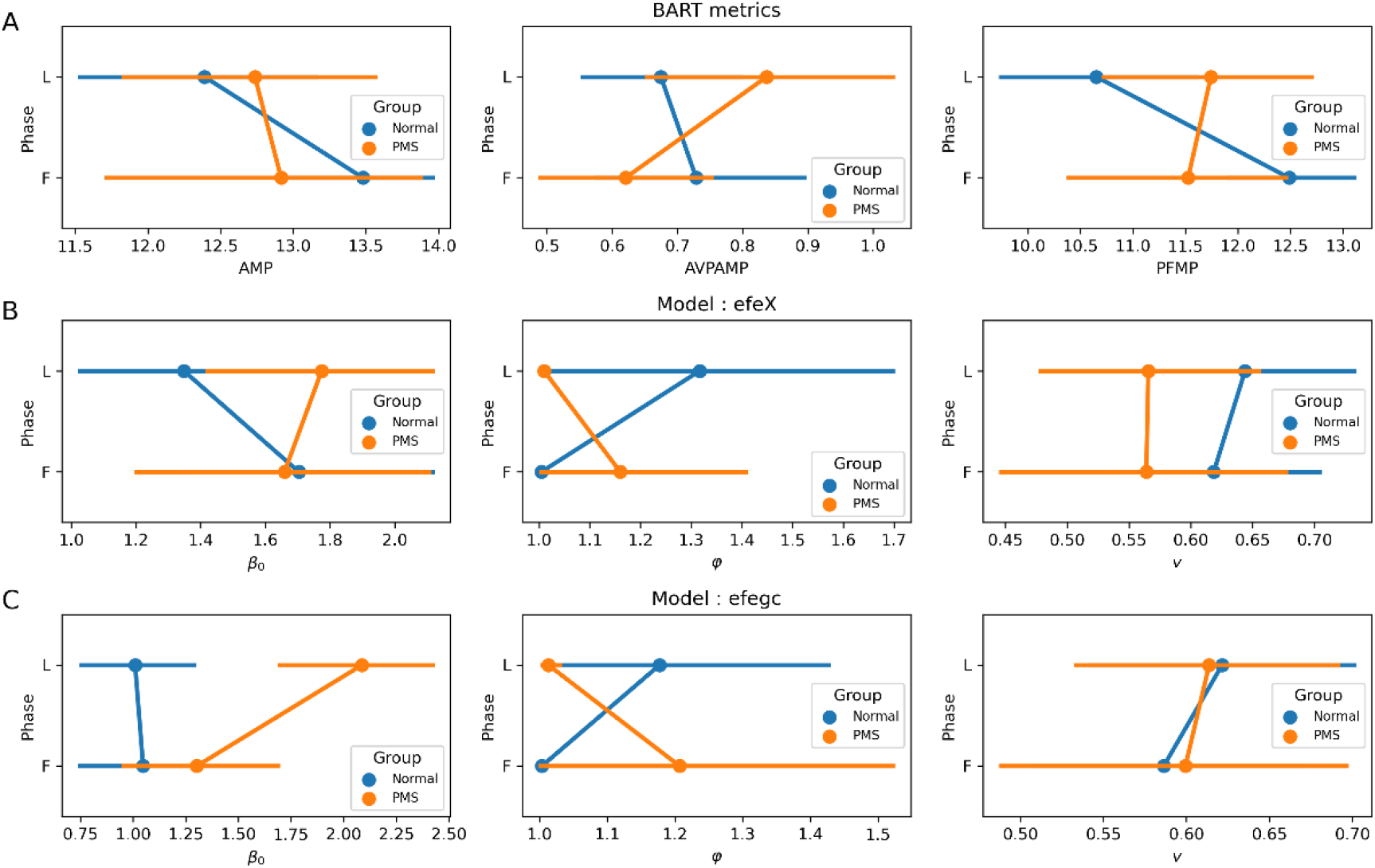
Comparison of BART behavioral metrics and computational model parameters across groups (Normal vs. PMS) and menstrual phases (Luteal [L] vs. Follicular [F]). (A) shows traditional BART metrics (AMP, AVPAMP, PFMP), (B) displays estimated parameters for the *efeX* model, and (C) shows estimated parameters for the *efegc* model. The parameter in the *efegc* model demonstrates a significant main effect of group, being consistently higher in the PMS group across both phases.

In the *efeX* model, no significant effect was found (Figure 5B and Supplementary Table 1). In the *efegc* model, the parameter *β*_0_ showed a significant main effect of group (F(1,151) = 17.27, p = .0001), with higher values in the PMS group. In addition, phase × group interaction was also significant (F(1,151) = 5.95, p = .016). Post-hoc comparisons revealed a significant effect of Phase within the PMS group (t = -2.83, p = .006), whereas no such difference was observed in the control group (t = .19, p = .853 in the control group). These results indicate that phase-related variations were specific to the PMS group.

## Discussion

In this study, we demonstrated that our novel AIF model effectively addresses and overcomes several core limitations of existing computational models for the BART. Previous models have typically treated sequential and prior evaluation strategies independently, or they have struggled to meaningfully capture uncertainty-driven information-seeking behavior. For instance, traditional models often rely on highly restricted parameter ranges where the variance component (intended to reflect uncertainty) becomes effectively negligible. Furthermore, continuous updating rules in prior models fail to reflect the discrete, step-like nature of human learning in specific task environments. By conceptualizing behavior as the minimization of EFE, our AIF model naturally integrates both pragmatic value (reward pursuit) and epistemic value (uncertainty reduction) into a single computational principle. This approach not only unifies initial pre-planned strategies with pump-by-pump sequential re-evaluations, but also accurately captures cognitive trajectories by modeling belief updates at discrete pump opportunities.

A critical feature of our proposed *efegc* model is the dynamic updating of policy precision, which operates in conjunction with EFE to explain complex risk-taking and gambling-like behaviors. Early in a trial, when the EFE of the stop policy is higher than that of the pump policy, successful pumps yield low VFE. This alignment between VFE and EFE increases policy precision, thereby amplifying the EFE-driven choice to continue pumping. Crucially, as the trial progresses and the risk of bursting grows, the EFE curves intersect, making the EFE of the pump policy higher than that of the stop policy—indicating that stopping is the rational choice. However, if a pump is successful at this highly uncertain stage, the unexpected nature of this success causes the directions of VFE and EFE to diverge, leading to a sudden decrease in policy precision. This reduction counteracts the rational influence of EFE, effectively suppressing the tendency to execute the stop action. This mechanistic interaction provides a compelling computational explanation for a common phenomenon observed in gambling: even when individuals have a pre-planned point at which they intend to stop and secure their winnings, a streak of consecutive rewards (analogous to successful pumps) dynamically lowers their policy precision, overriding their rational evaluation and driving them to continue gambling beyond their originally intended limit.

In the analysis in PMS group, although the estimated parameter values from the *efegc* and *efeX* models showed high correlations—*β*_0_ (*r* = .472,*p* < .0001),*φ*(*r* = .650,*p* < .0001), and *υ* (*r* = .743, *p* < .0001)—the two models yielded different results. We confirmed that these differences stem from the distinct behavioral characteristics each parameter reflects, as evidenced by their correlations with the BART metrics. Notably, in the *efeX* model, *β*_0_ is significantly correlated with AMP and PFMP, but not with AVPAMP (Supplementary Table 3). In contrast, the *efegc* model revealed a positive correlation between *β*_0_ and AVPAMP. A higher AVPAMP indicates a lack of overall consistency in pump counts, implying that sequential evaluation exerts a stronger influence than prior evaluation during decision-making. Furthermore, no significant group differences were observed in the parameter *υ*, which is involved in belief updating. While *υ* showed a positive correlation with AVPAMP, it also showed a negative correlation with PFMP without positive correlation with AMP. This high correlation with PFMP suggests that the high *υ* reflects a sensitivity to unexpected observations from preceding trials (e.g., a marked reduction in pumps immediately following an explosion) rather than global behavioral inconsistency. *β*_0_ also showed a negative correlation with PFMP, but it also showed a significant negative correlation with AMP, which naturally increases the PFMP. After adjusting for AMP, the correlation between *β*_0_ and this index disappeared in the *efegc* model (PFMPAMP in Supplementary Table 3), suggesting that the initial correlation between *β*_0_ and PFMP was likely confounded by AMP. Thus, the positive correlation between *β*_0_ and AVPAMP suggests that it is independent of any failure or deficit in learning. This is further supported by the observation that PFMP was not specifically lower in the PMS group; had learning been impaired by a high decay term driven by elevated *υ*, a significant reduction in PFMP would be expected. Therefore, independent of the learning process, the elevated *β*_0_ in the PMS group points to a lack of behavioral consistency during the action phase itself. This reflects a form of impulsivity characterized by on-the-spot choices rather than planning and executing long-term actions. The result that this difference was observed only in the PMS group suggests that increase of *β*_0_ may serve as a marker reflecting the worsening of mood or cognitive symptoms across the menstrual cycle in individuals with PMS. However, because the data for the two phases were not obtained through repeated within-subject measurements, the current sample size is insufficient to draw meaningful conclusions regarding the phase effect. Therefore, future studies should validate these phase-related findings using repeated-measures datasets.

## Conclusion

We introduce a novel computational model based on AIF that effectively captures the complex psychological dynamics of decision-making and risk-taking in BART task. In addition, by modeling belief updates at discrete pump opportunities and incorporating dynamic policy precision modulated by variational free energy, the proposed *efegc* model unifies sequential and prior evaluation strategies and statistically outperforms existing conventional models. Applying this framework to women with PMS provided deeper mechanistic insights into impulsivity that traditional behavioral metrics could not capture. Specifically, our model revealed that individuals with PMS exhibit significantly higher values, indicating that their impulsivity is driven by on-the-spot behavioral inconsistency during the action phase, rather than a fundamental learning deficit. Although the phase-dependent effects require further validation through repeated-measures designs, this research establishes AIF as a powerful tool in computational psychiatry, offering a more granular and clinically relevant approach to understanding the cognitive mechanisms underlying impulsive behaviors.

## Data and code availability

The data underlying this article will be shared upon reasonable request to the corresponding author. The code will be available by requesting to the first author.

## Acknowledgments

This study was supported by a Medical Scientist Training Program from the Ministry of Science & ICT of Korea, grant of the Korea Health Technology R&D Project through the Korea Health Industry Development Institute (KHIDI) [RS-2024-00440131] and the Brain Research Program through the National Research Foundation of Korea (NRF) funded by the Ministry of Science & ICT [RS-2025-00562004].

## Supplementary materials

Supplementary.pdf

