## Supplementary Figure 1-3, Table 1-3 for "Policy precision reveals action-phase impulsivity in women with premenstrual syndrome during risk-taking"

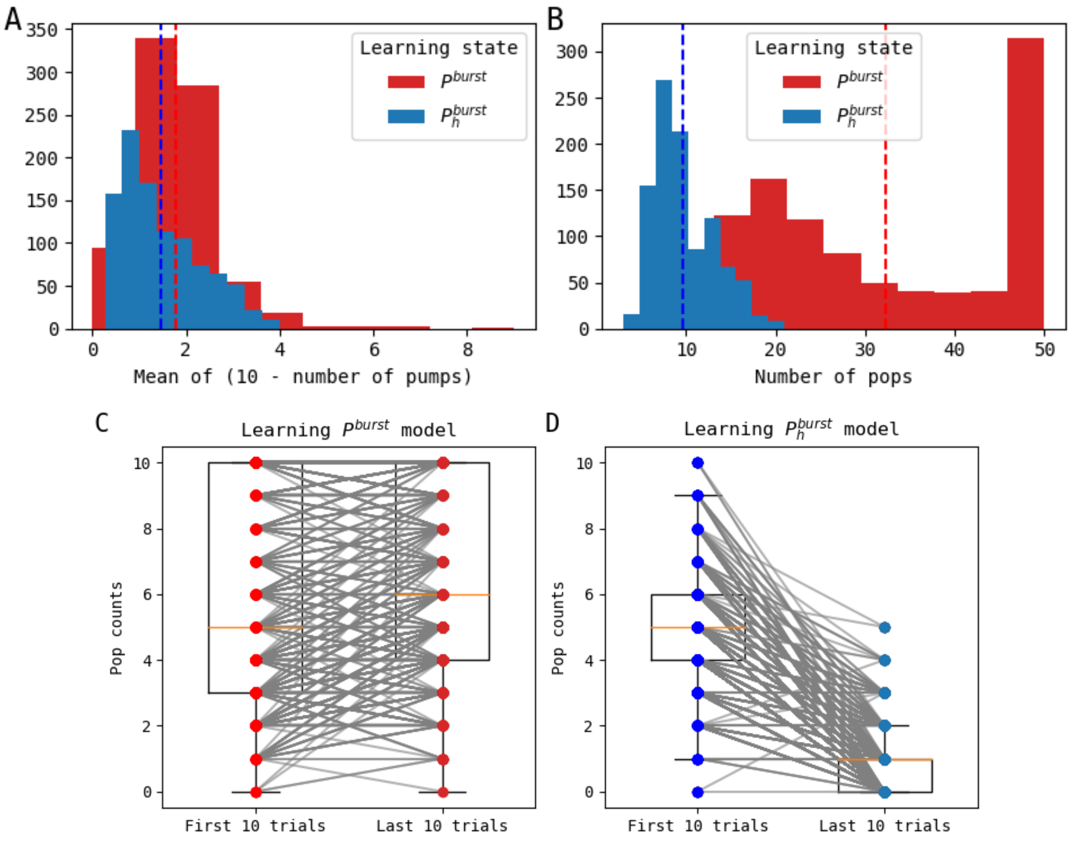


Figure 1. Simulation results of a deterministic task environment where the balloon invariably bursts at the 10th pump. Model parameters were randomly sampled from a uniform distribution, and 1,000 simulations were generated per model. (A) Histogram of the mean difference between the maximum possible pumps (10) and the actual number of pumps in successful (non-burst) trials. (B) Histogram of total pop counts per model. The conventional learning model ($p^{burst}$) exhibits a significantly higher frequency of trials where all 50 balloons burst compared to the novel learning model ($\boldsymbol{p_h^{burst}}$) ($t$ = 51.448, $p$ < .0001). (C, D) Comparison of pop counts between the first 10 trials and the last 10 trials to assess learning progression. The novel model (D) demonstrates a significant decrease in pops over time (t = -16.013, p < .0001), indicating successful learning of the deterministic rule, whereas the conventional model (C) fails to adapt, rather showing an increase in pop counts (t = 73.87, p < .0001).

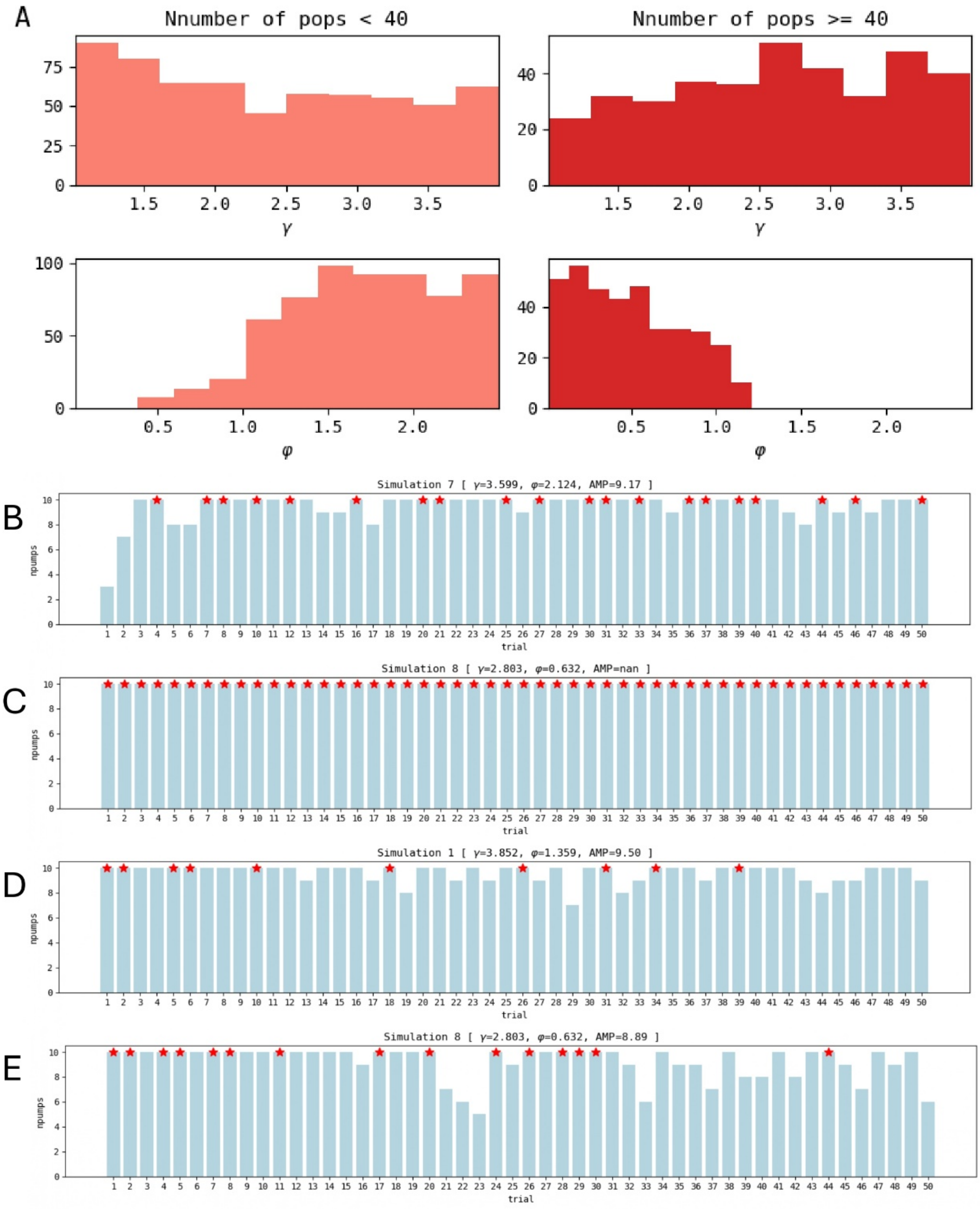


Figure 2. Parameter distributions and example simulation plots for the conventional learning models ($p^{burst}$ model in Figure 1). (A) Histograms of parameter distributions ($\gamma$ and $\varphi$), categorized by simulations with total pop counts being lower than 40 and over 40. The $\gamma$ parameter is inverse of $\beta_0$ and shows no clear distributional difference, but a loss aversiveness ($\varphi$) value of less than 1 consistently results in pops across nearly all trials. (B) An example simulation plot for the conventional learning model with pop counts lower than 40 and even in this, the frequency of successful trials remains unchanged between the early and late phases of the task. (C) An example simulation plot for the model with pop counts over 40. (D, E) An example simulation plot generated by the novel learning model with both high and low $\varphi$ values.

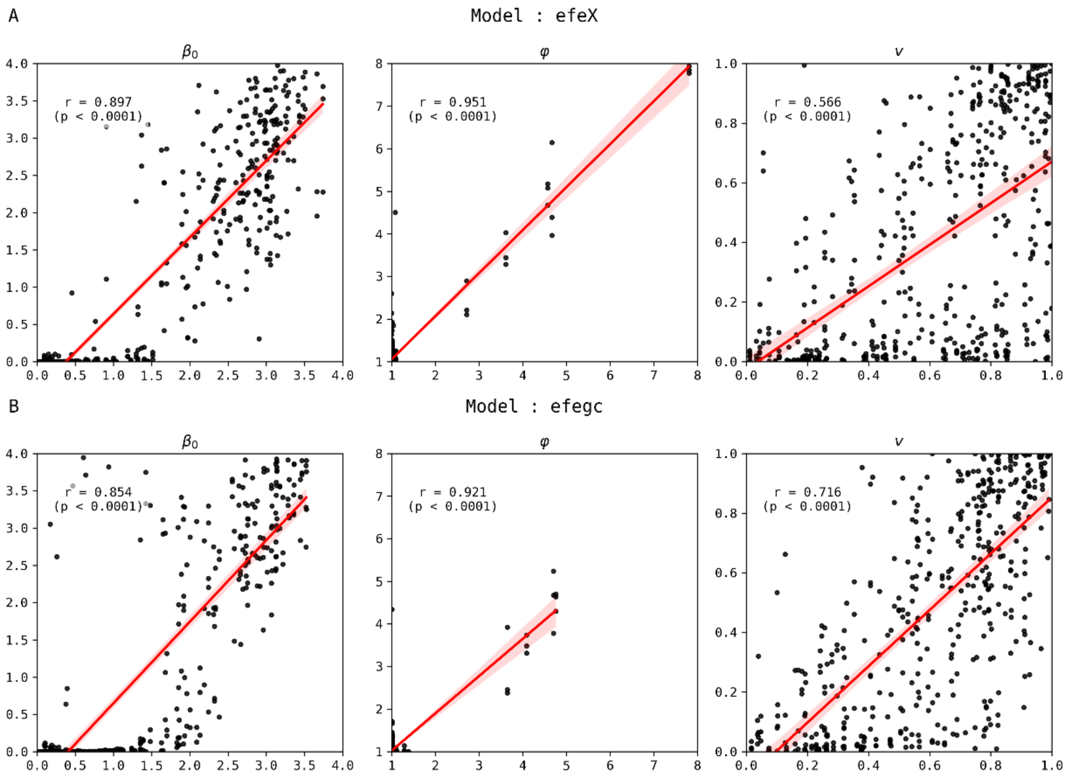


Figure 3. Parameter recovery results for both active inference models (*efeX* and *efegc*). The x-axis represents the true parameter values, and the y-axis represents the recovered parameter values. The ranges of the x- and y-axes correspond to the ranges used to estimate each parameter.

**Table 1. Two-Way ANOVA results for model parameters and BART behavioral metrics across menstrual phase and group**

| Model | Parameter | Phase (F, p) | Group (F, p) | Phase × Group (F, p) |
|---|---|---|---|---|
| *efeX* | $\beta_0$ | .57 (.450) | 1.10 (.296) | 1.38 (.242) |
| | $\varphi$ | .98 (.324) | .63 (.430) | 3.81 (.053) |
| | $v$ | .06 (.809) | 1.87 (.174) | .06 (.811) |
| *efegc* | $\beta_0$ | **4.245 (.041)*** | **17.27 (.0001)**** | **5.95 (.016)*** |
| | $\varphi$ | .06 (.801) | .00 (.952) | **4.41, (.038)*** |
| | $v$ | .37 (.544) | .00 (.967) | .06 (.815) |
| **BART metrics** | | **Phase (F, p)** | **Group (F, p)** | **Phase × Group (F, p)** |
| AMP | | 3.06 (.083) | 0.03 (.862) | 1.19 (0.277) |
| AVPAMP | | 0.58 (.448) | 0.22 (.641) | 2.83 (0.094) |
| PFMP | | **4.76 (.031)*** | 0.10 (.750) | **5.08 (0.026)*** |

**Table 2. Pearson correlation results between model parameters and psychiatric symptom scales**

| Model | Parameter | Symptom scales | | | | | | | | | |
|---|---|---|---|---|---|---|---|---|---|---|---|
| | | PSST | DRSP | RAS | RSES | PHQ | GAD | STAI2 | MAIA | BIS | DERS |
| *efeX* | $\beta_0$ | - | - | -0.167 (0.037) | -0.163 (0.042) | - | -- | - | - | - | - |
| | $\varphi$ | - | - | -0.170 (0.033) | -0.196 (0.014) | - | 0.213 (0.007) | -0.179 (0.025) | -0.179 (0.025) | - | - |
| | $v$ | - | - | - | - | - | - | - | - | - | - |
| *efegc* | $\beta_0$ | **0.279 (0.0004) **** | 0.25 (0.002) | -0.207 (0.009) | - | - | 0.189 (0.018) | 0.247 (0.002) | - | - | 0.211 (0.008) |
| | $\varphi$ | - | 0.191 (0.016) | -0.271 (0.001) | -0.272 (0.001) | 0.224 (0.005) | **0.367 (<.0001) **** | 0.213 (0.007) | -0.243 (0.002) | 0.199 (0.012) | 0.235 (0.003) |
| | $v$ | - | - | - | - | - | - | - | - | - | - |

**Table 3. Pearson correlation results between model parameters and BART behavioral metrics**

| Model | Parameter | BART metric | | | |
|---|---|---|---|---|---|
| | | AMP | AVPAMP | PFMP | PFMPAMP |
| *efeX* | $\beta_0$ | -0.299 (0.0001) | 0.101 (0.207) | -0.226 (0.004) | 0.093 (0.245) |
| | $\varphi$ | -0.489 (<.0001) | -0.013 (0.876) | -0.443 (<.0001) | -0.032 (0.691) |
| | $v$ | -0.103 (0.197) | 0.245 (0.002) | -0.192 (0.016) | -0.114 (0.155) |
| *efegc* | $\beta_0$ | -0.324 (<.0001) | 0.207 (0.009) | -0.181 (0.023) | 0.129 (0.108) |
| | $\varphi$ | -0.629 (<.0001) | -0.079 (0.325) | -0.488 (<.0001) | 0.037 (0.647) |
| | $v$ | -0.055 (0.495) | 0.416 (<.0001) | -0.223 (0.005) | -0.159 (0.046) |
| *Par4* | $\tau$ | 0.032 (0.688) | -0.063 (0.435) | -0.021 (0.79) | -0.116 (0.147) |
| | $rt$ | 0.363 (<.0001) | -0.170 (0.034) | -0.127 (0.114) | -0.961 (<.0001) |
| | $\eta$ | -0.127 (0.113) | 0.236 (0.003) | -0.156 (0.051) | -0.04 (0.618) |
| | $\phi$ | 0.229 (0.004) | 0.051 (0.527) | 0.175 (0.028) | 0.014 (0.863) |
| *EWMV* | $\tau$ | -0.560 (<.0001) | -0.06 (0.458) | -0.487 (<.0001) | -0.025 (0.76) |
| | $la$ | -0.500 (<.0001) | -0.114 (0.154) | -0.403 (<.0001) | -0.001 (0.987) |
| | $\eta$ | 0.440 (<.0001) | -0.146 (0.068) | 0.378 (<.0001) | 0.121 (0.131) |
| | $\phi$ | -0.149 (0.063) | 0.335 (<.0001) | -0.133 (0.097) | -0.06 (0.453) |
| | $rp$ | 0.443 (<.0001) | 0.115 (0.153) | 0.351 (<.0001) | -0.001 (0.988) |
